# On stability of the Darwinian dynamics

**DOI:** 10.1101/2021.10.26.465938

**Authors:** Mohammadreza Satouri, Jafar Rezaei, Kateřina Staňková

**Affiliations:** Faculty of Technology, Policy and Management, Delft University of Technology, Delft, The Netherlands

**Keywords:** Darwinian dynamics, Lyapunov stability, mathematical oncology

## Abstract

Here we analyze Darwinian dynamics of cancer introduced in [1], extended by including a competition matrix, and evaluate (i) when the eco-evolutionary equilibrium is positive and (ii) when the eco-evolutionary equilibrium is asymptotically stable.

## 1 Introduction

Game-theoretical models help us with understanding cancer and its treatment [2, 3]. In [1], one such a model, describing eco-evolutinary dynamics of two cancer cell subtypes, one being completely treatment-sensitive and another one, evolving treatment-induced resistance, were considered. Different treatment regimens, which could be more effective than commonly used maximum tolerable dose, were analyzed. The interaction between different cancer subtypes happened only through common carrying capacity. Here we investigate whether in a model of [1] with cost of resistance in the intrinsic growth rate, expanded by a competition matrix, we can achieve stable and viable eco-evolutionary equilibria.

According to the Hartman-Grobman theorem, by linearizing a nonlinear dynamic around its equilibrium points, we can judge stability of equilibria in a vicinity of those points (local stability) [4]. If at least one of the eigenvalues of the Jacobian matrix corresponding to a particular equilibrium lies on the imaginary axis, no judgment can be made on the stability of that equilibrium and other methods, such as center manifold theory [5], should be utilized. Moreover, due to the linearization, no information can be obtained about the basin of attraction and global stability cannot be discussed. That is why we adopt the Lyapunov theory to analyze stability of the eco-evolutionary equilibria, to see under which conditions they are globally asymptotically stable.

## 2 The model

We model the evolution of resistance leading to treatment failure using ordinary differential equations for a polymorphic tumor cell population. The entire tumor cell population is comprised of two distinct subpopulations: sensitive and resistant cells (populations *x*_*S*_(*t*) and *x*_*R*_(*t*), respectively). In this model, only the resistant cell subpopulation has the capacity to evolve resistance as a quantitative trait *u*_*R*_(*t*) ∈ ℝ_+_. We assume that the tumor subpopulations grow logistically and are suppressed by the presence of therapy and natural cell turnover. The model describes Darwinian dynamics of cancer in response to treatment, with a fitness-generating function, “G-function” [6]. A G-function describes how the fitness of a focal cancer cell using a strategy *v*(*t*) in the population is influenced by the environment and by the strategies and population sizes of the resident subtypes. The set of strategies present in the tumor are represented by **u**(*t*). The population size of cells with a particular strategy is indicated by **x**(*t*). In the polymorphic context, the vector **u**(*t*) = (*u*_*R*_(*t*), *u*_*S*_(*t*))^*T*^ encompasses the strategy for resistant and sensitive cells and **x**(*t*) = (*x*_*R*_(*t*), *x*_*S*_(*t*))^*T*^ their population sizes. We assume that the physician applies a treatment dose *m*(*t*) ∈ [0, 1] at time *t* ≥ 0, where *m*(*t*) = 0 and *m*(*t*) = 1 correspond to no dose and MTD at time *t*, respectively. For simplicity, the drug is assumed to be maximally effective at MTD. The efficacy of the drug is reduced by a focal cell’s resistance strategy *v*, innate drug immunity *k*, and the benefit *b* of the resistance trait in reducing therapy efficacy. The G-function is used to derive the evolutionary dynamics that describe how the resident strategies (i.e. subtypes) of the tumor change with time. Following Fisher’s fundamental theorem of natural selection, the resistance strategies change in the direction of the fitness gradient 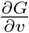 with respect to the fitness of a rare mutant *v*(*t*) [7]. This derivative is then evaluated at the current resident strategies **u**(*t*), giving an equation defining the evolutionary dynamics for each resident strategy [6]. The rate at which the strategies change is scaled by an evolutionary speed term *σ*. In our model, large values of evolutionary speed *σ* correspond to enhanced phenotypic variance which could result from increased genetic variance or phenotypic plasticity. Innate immunity *k* suggests that prior to drug exposure cells possess a mechanism that inhibits the potency of treatment. This parameter is the only value that reduces drug efficacy for the sensitive population in our polymorphic model as the sensitive cells cannot evolve resistance. Treatment efficacy is further diminished by the magnitude of the benefit *b* of the resistance strategy for the monomorphic population and the resistant population in the polymorphic model. For a general introduction to our modelling framework.

The eco-evolutionary dynamics of our polymorphic population can be then described for *i* ∈ {*R, S*} as follows:

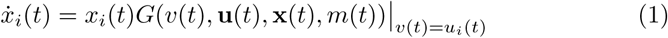

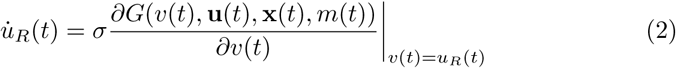

Here the fitness-generating function is defined as

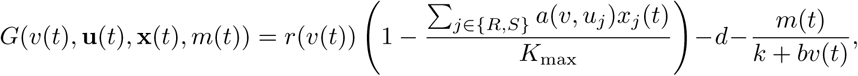

with *a*(*u*_*i*_, *u*_*j*_) = *α*_*ij*_, *α*_*ij*_ ∈ [0, 1] being a competition effect of type *j* on type *i, r*(*v*(*t*)) = *r*_max_ *e*^−*g v*(*t*)^, and we assume that *α*_*ii*_ = 1 and *u*_*S*_(*t*) = 0 for all *t*. Using notation

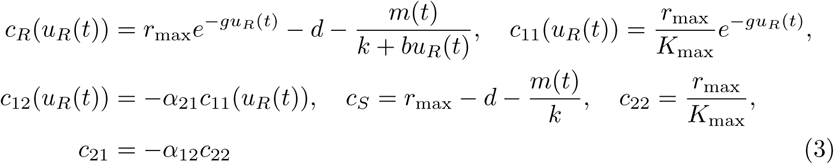

we can rewrite dynamics (1)-(2) into the following form:

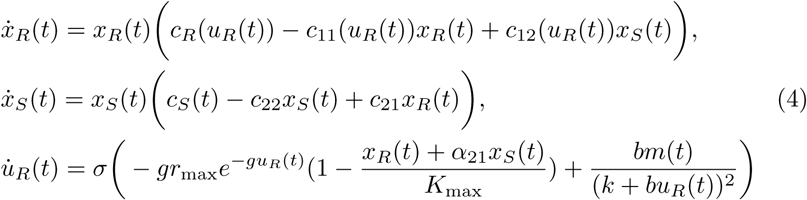

## 3 Stability of interior eco-evolutionary eqilibria via Lyapunov approach

Here we focus on interior equilibria. At first, the necessary conditions for having interior equilibria are obtained, and then the stability of this equilibria is investigated. In the sequel, the time dependence is dropped for simplicity.

The equilibriums of dynamics (4) are

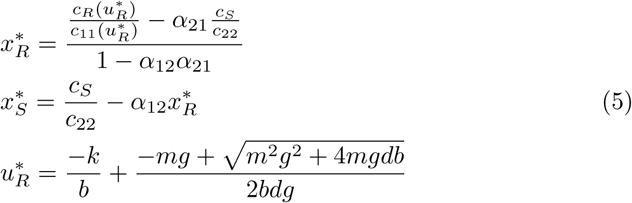

According to (5), conditions for having interior eco-evolutionary equilibrium points are as follows:

1. *α*_*12*_ *α*_*12*_ *< 1*
2. 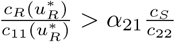
3. 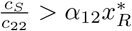
4. *K*^*2*^*dg + kmg < mb*

From condition 3, one can obtain:

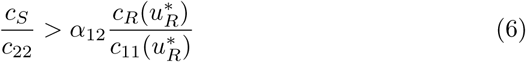

If 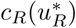 and *c*_*S*_ have different signs, then from Condition 3, and (6), it is concluded that one of *α*_12_ or *α*_21_ should be negative. Consequently, 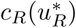 and *c*_*S*_ should have same signs.

Now, the asymptotic stability of the eco-evolutionary equilibrium points is investigated. Augmenting cancer cells’ dynamics with the dynamics of resistant cells’ strategies, a new variable ***ξ*** = (*x*_*R*_, *x*_*S*_, *u*_*R*_)^*T*^ is introduced.

### Theorem 1.

*Considering dynamics* (4), *under assumptions 1-4, the interior eco-evolutionary equilibrium is globally asymptotically stable*.

*Proof*. For this purpose, the following candidate Lyapunov function is introduced.

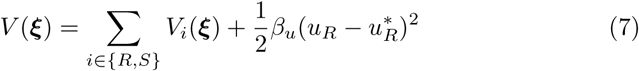

in which

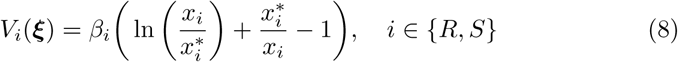

### Remark 1.

*The function* ln 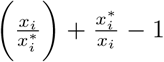 *is increasing for* 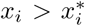, *and is decreasing for* 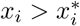. *Also, it is zero at* 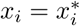.

### Remark 2.

*As can be seen, the candidate Lyapunov function* (7) *is radially unbounded*.

Time derivative of the candidate Lyapunov function is:

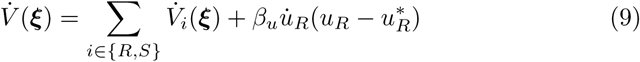

where

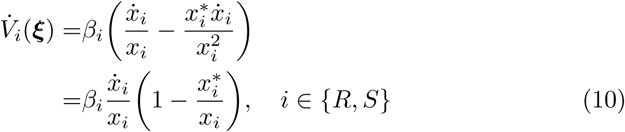

Substituting dynamics (4) in (10), yields

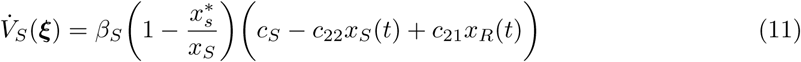

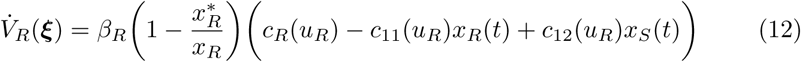

At the equilibrium point, one can obtain

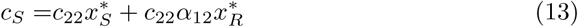

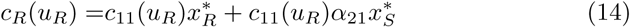

By substituting (13) in (11) and using this fact that *c*_21_ = −*α*_12_*c*_22_ we have

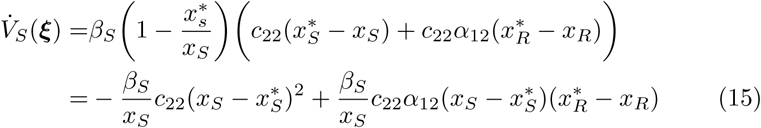

With a similar approach one can obtain

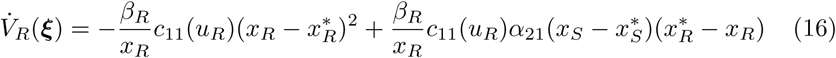

According to (15), and (16), if 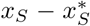 and 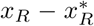 have a same sign, then both of 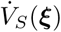 and 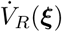 become negative definite. In this case, by increasing *β*_*S*_ and *β*_*R*_, and decreasing *β*_*u*_, regardless of the sign of 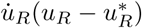, the time derivative of the Lyapunov function (9) becomes negative. In the sequel, we will show via contradiction that for the remaining cases, 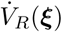 and 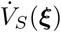 cannot be simultaneously positive.

We assume that 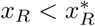 and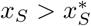, and rewrite (15) and (16) as follows:

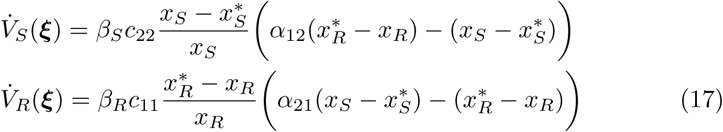

If we assume that 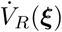 and 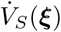 are positive simultaneously, it yields

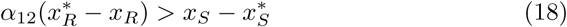

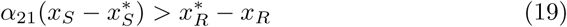

By multiplying (18) with *α*_21_ we have

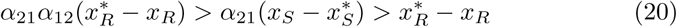

This means that *α*_21_*α*_12_ *>* 1 which contradicts with condition 1. As a result, 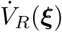 and 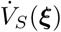 cannot be simultaneously positive. This fact can be proved in a similar way for the case 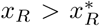 and 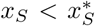. Thus, always at least one of the 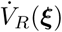 or 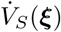 is negative. In this case, we increase the coefficient of the 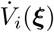 which is negative, and decrease the other coefficients, and with this approach, the time derivative of the candidate Lyapunov function (7) is always negative for all the interior equilibrium points.

## 4 Conclusion

The conditions for having an interior eco-evolutionary equilibrium of cancer Darwinian dynamics was obtained and the asymptotic stability of these equilibria was proved.

## Acknowledgements

This research is supported by European Union’s Horizon 2020 research and innovation programs under the Marie Skłodowska-Curie grant under the grant agreement number 955708.

